# Endothelial cell Nrf2 controls neuroinflammation following a systemic insult

**DOI:** 10.1101/2024.10.21.619507

**Authors:** Haoyu Zou, Tom Leah, Zhuochun Huang, Xin He, Eleonora Mameli, Andrea Caporali, Owen Dando, Jing Qiu

## Abstract

Systemic inflammation can lead to neuroinflammation with acute consequences such as delirium and long-lasting deleterious effects including cognitive decline and the exacerbation of neurodegenerative disease progression. Here we show that transcription factor Nrf2 controls brain endothelial cell homeostasis and barrier strength. We found that peripheral inflammation caused infiltration of macrophages, microglial activation and inflammatory reactive astrogliosis, all of which could be prevented by RTA-404, an activator of the transcription factor Nrf2 and close structural relative of the recently FDA-approved Nrf2 activator RTA-408 (Omaveloxolone). To identify the key cellular mediator(s), we generated an endothelial cell-specific Nrf2 knockout mouse. Strikingly, the effects of RTA-404 on brain endothelial activation and downstream neuroinflammatory events was abolished by endothelial cell-specific Nrf2 deletion. This places endothelial cell Nrf2 as a peripherally accessible therapeutic target to reduce the CNS-adverse consequences of systemic inflammation.

## Introduction

Systemic inflammation is a major cause of neuroinflammation, key markers of which include infiltration of circulating leukocytes into the brain parenchyma across a compromised blood brain barrier (BBB), as well as microglial activation and inflammatory reactive astrogliosis^1–6^. Neuroinflammation can lead acutely to delirium, and chronically to exacerbation of neurodegenerative disease progression including vascular dementia and Alzheimer’s disease^7–11^. Strategies to prevent CNS-adverse consequences of systemic inflammation require a better knowledge of the key pathways that maintain the immune-privileged status of the CNS. Brain endothelial cells (BECs) are an integral component of the BBB, and therefore an attractive proposition as a target for immune system-to-brain signalling^12–14^.

The transcription factor Nrf2 (encoded by the *Nfe2l2* gene) is a widely-expressed stress-responsive master regulator of genes involved in important aspects of homeostatic physiology including the prevention of redox, metabolic and inflammatory deregulation^15–17^. Under normal conditions Nrf2 is targeted for ubiquitin-mediated degradation by Keap1, but in response to stress, this interaction is inhibited, leading to Nrf2 accumulation in the nucleus where it regulates the expression of phase II and antioxidant defence enzymes containing antioxidant response elements (AREs) in their promoter^15,17^, including some of the canonical Nrf2 target genes heme oxygenase-1 (HO)^18^, NAD(P)H:quinine oxidoreductase-1 (NQO1)^19^, glutamate–cysteine ligase modifier subunit (GCLM)^20^, and thioredoxin reductase 1 (TXNRD1)^21^, which protect the cells against oxidative and electrophilic stress. In addition, it has been suggested that Nrf2 can control over 1000 other genes with functions ranging from antioxidant defence, anti-inflammation, detoxification to proteostasis and metabolism^22^. Several previous studies have shown the beneficial roles of Nrf2 in endothelial cells (ECs)’ development and homeostasis. For example, global and EC-specific deletion of Nrf2 demonstrated a critical role of Nrf2 in angiogenesis during vascular development^23^. Nrf2 protected vascular ECs from oxidative stress and inflammation and enhanced EC integrity via activating Nrf2 target genes e.g. HO and NQO1^24,25^. Global Nrf2 deletion increased susceptibility to BBB damage in animal disease models^26–29^.

Cyano-3,12-Dioxooleana-1,9-Dien-28-Oic acid (CDDO) and its analogues, e.g. RTA-404 (CDDO-TFEA) and RTA-408 (Omaveloxolone), are a class of synthetic oleanane triterpenoid compounds and by far the most potent small molecule Nrf2 activators via disrupting KEAP1-Nrf2 interactions, leading to stabilization and rapid nuclear translocation of Nrf2 and transcription of Nrf2 target genes^28,30,31^. For the last two decades, the triterpenoid compounds have exhibited a broad range of applications with their anti-inflammatory and antioxidative properties in animal models and clinical trials^32,33^. Compared to other triterpenoid compounds, RTA-404 has greater capacity to cross the BBB and has shown neuroprotective effects in animal models of neurodegenerative diseases, including Autoimmune encephalomyelitis^34^, Huntington′s disease^35^, and Amyotrophic lateral sclerosis^36^. RTA-408 was recently approved by the FDA for treating Friedreich’s Ataxia, a genetic neurodegenerative disease associated with oxidative stress^37^. This provides proof-of-concept that Nrf2 can be safely activated by the triterpenoid compounds in humans. In the brain, Nrf2 is highly expressed in BECs, microglia and astrocytes, with low expression in neurons^38–40^. However, cell-type specific roles of Nrf2 in the brain are not well understood, partly due to a historical reliance on global Nrf2 knockout mice to probe the roles of Nrf2^41^. This knowledge gap is important to fill because cell type-specific roles of Nrf2 are likely to be influenced by the function of that cell type, and from a translational point of view one ideally needs to know the locus of action of any Nrf2-targeting therapeutic in order to optimise bioavailability. Previously, our work has focussed on the impact of RTA-404 on astrocytes. We demonstrated that *in vitro* RTA-404 triggers a neuro-protective response in astrocytes via a Nrf2-dependent transcriptional programme^42^. However, the effects of RTA-404 and other CDDO analogues on BECs were unknown.

## Results

### The Nrf2 activator RTA-404 promotes brain endothelial cell barrier strength

To test the influences of RTA-404 on BECs, we assessed the effects of RTA-404 on human BEC barrier strength by measuring trans-endothelial electrical resistance (TEER) *in vitro*. We found that RTA-404 induced an increase in BEC TEER (Fig. 1A). We also performed RNA-seq analysis to look at the effects of RTA-404 on BEC transcriptome at basal conditions. RTA-404 significantly activated a number of prototypical Nrf2 target genes, e.g. *HMOX1, NQO1, TXNRD1, FTL, FTH1 and GCLM* (Fig. 1B). In addition, the expression of several genes involved in tight junctions and adherent junctions, e.g. *OCLN*, *JHY, ANTXR1, MYZAP* and *AFDN* were significantly upregulated with RTA-404 (Fig.1B). The Ingenuity Pathway Analysis (IPA) of genes significantly altered suggested that pathways, e.g. NRF2 regulating anti-oxidant/detoxification enzymes, Iron uptake and transport, NRF2 regulating pentose phosphate pathway, iNOS signalling and IL10 signalling were activated by RTA-404 (Fig. 1C and Table S1). The activation of Nrf2 with RTA-404 in human BECs was also investigated with immunocytochemistry showing an increased in protein expression of NRF2, NRF2 target genes HMOX1 and NQO1, as well as tight junction protein ZO1 (Fig. 1D).

**Fig.1.**
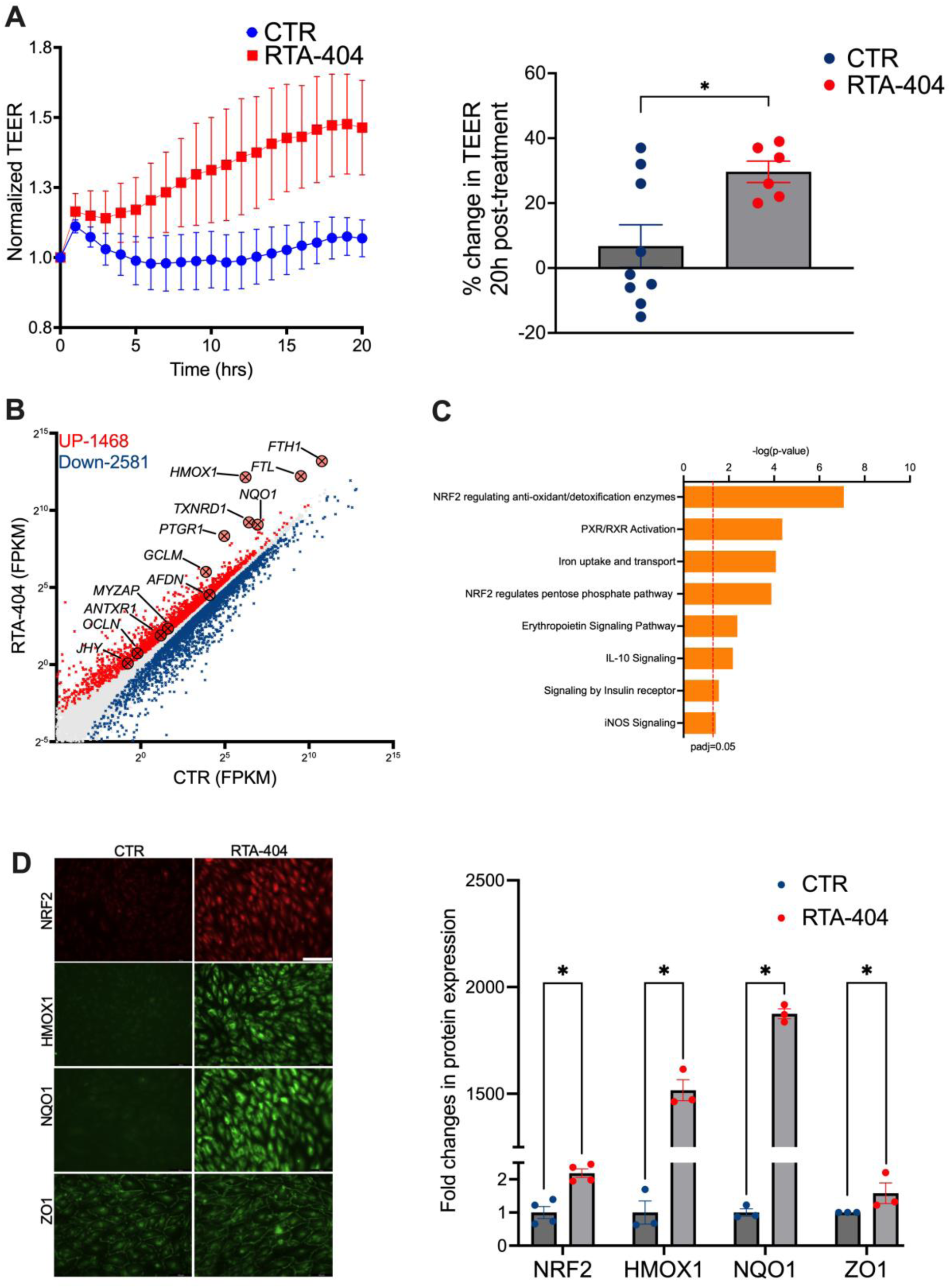
Nrf2 activator RTA-404 promotes brain endothelial barrier strength. **(A)** Human BEC cultures were treated with ±RTA-404 and transepithelial/transendothelial electrical resistance (TEER) was measured for 20 hrs. *p=0.0136, One-way ANOVA plus Sidak post hoc (n=8-9). **(B)** Human BEC cultures were treated with ±RTA-404 for 24h, RNA was extracted and RNAseq was performed. Scatterplot was generated for genes with average expression >0.1 FPKM across the data sets. Highlighted with red and blue crosses are the genes whose expression are significantly increased or decreased respectively (*DESeq2 P*_adj<0.05, n=3). ‘N’ refers here and throughout as an independent biological replicate (i.e. an independent culture or a mouse). **(C)** IPA analysis identifying activated pathways in human BEC culture by treated with RTA-404 (RTA-404 vs. DMSO Control). Significantly DEGs (*DESeq2 P*_adj<0.05, |Log_2_FC|>1, n=3) with BEC transcriptome as reference dataset were analyzed to calculate the p-value of overlap and z-score of overall activation/inhibition states of individual pathways. The significant activated (p<0.05, z-score >2) pathways are shown in the bar chart. **(D)** Immunocytochemistry showing RTA-404 activates Nrf2 nuclear translocation and upregulates the protein expression of Nrf2 target genes *HMOX1* and *NQO1*, as well as tight junction protein ZO1. From left to right p=0.0008, 0.0005, 0.0002, and 0.05, n=3-4, unpaired t-test. Scale bar = 100 um.

### Endothelial cell-specific knockout of Nrf2 impaired BEC homeostasis and reduced endothelial cell barrier strength

We wanted to further investigate the role of Nrf2 in the BECs themselves. We used *Nefe2l2^fl/fl^* mice which possess *loxP* sites on either side of exon 5 of the *Nefe2l2* gene that encodes the DNA binding domain Nrf2 protein. *Nefe2l2^fl/fl^* mice were crossed onto a line, *Cdh5^CreERT^*^28^ which expresses tamoxifen-inducible CreERT specifically in endothelial cells, to generate an inducible-conditional EC-specific knockout line (*Cdh5^CreERT^*^2^*:Nfe2l2^fl/fl^*, hereafter *Nfe2l2*^ENDO^). Sorting of brain cells by FACS followed by qPCR confirmed successful deletion of Nrf2 exon 5 in BECs, without affecting Nrf2 levels in astrocytes, microglia or oligodendrocytes (Fig. 2A). When cultured BECs isolated from *Nfe2l2*^ENDO^ mice, we observed a significant decrease in endothelial barrier strength compared to those from control *Nefe2l2^fl/fl^* mice (Fig. 2B), suggesting that Nrf2 is important in maintaining endothelial barrier strength. RNA-seq analysis showed that Nrf2-deficient BECs exhibited a down-regulation of several known Nrf2 target genes (e.g. *Txnrd1* and *Gstm1*, Fig. 2C), as well as genes involved in forming tight junctions, e.g. *Jam3*, although the modest magnitude of changes observed suggested that basal Nrf2 activity in BECs may be relatively low. Changes caused by Nrf2-deficiency in BECs were also observed at the protein levels, assessed by mass spectrometry analysis (Fig. 2D), including a down-regulation of Nrf2 target genes such as *Txnrd1*, *Prdx1* and *Prdx3*.

**Fig.2.**
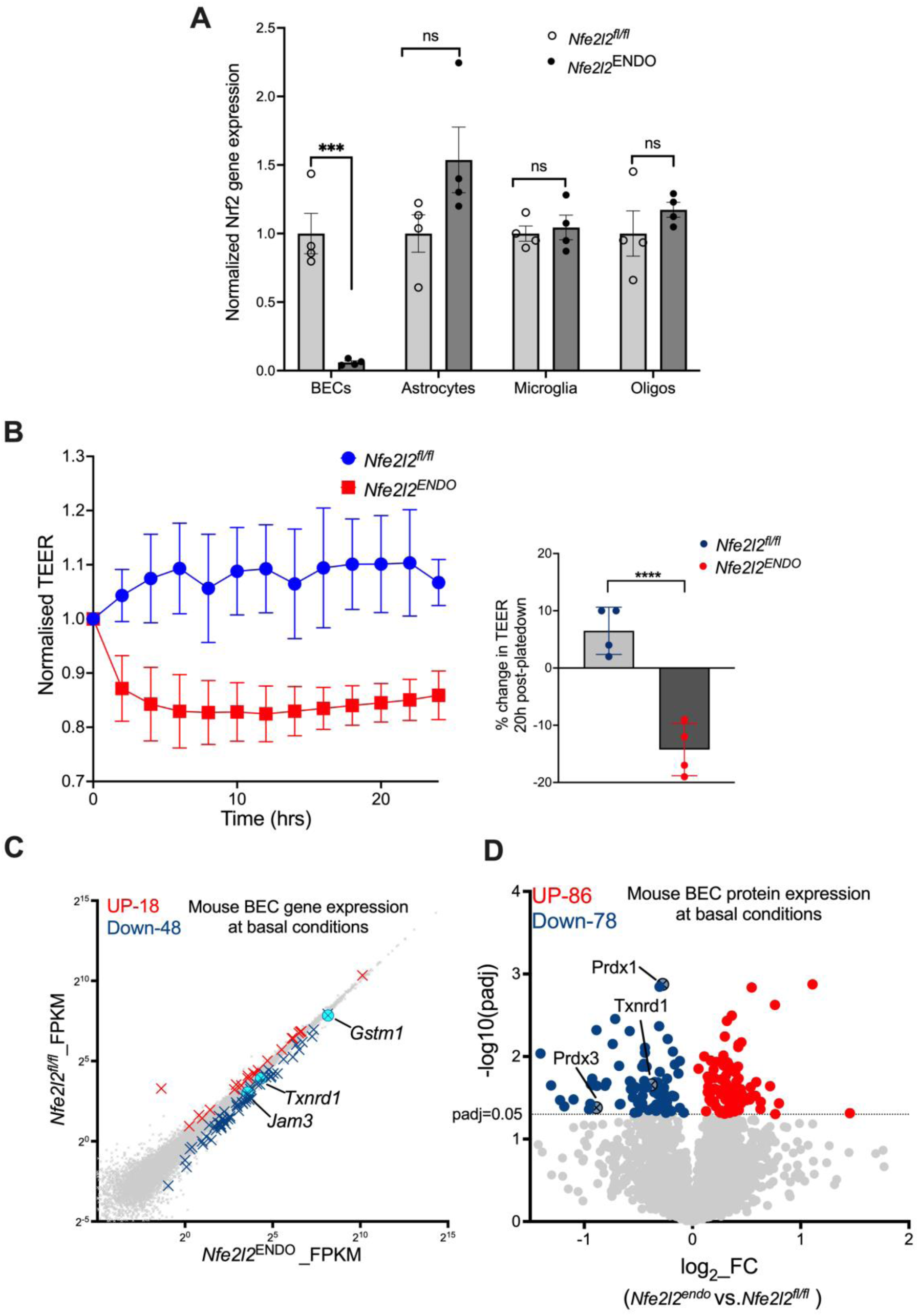
Endothelial cell-specific knockout of Nrf2 impaired BEC homeostasis and reduced endothelial barrier strength. **(A)** qRT-PCR confirms gene expression of exon 5 of Nrf2 is specifically deleted in BECs, but not in astrocytes, microglia or oligodendrocytes in *Nfe2l2*^ENDO^ mice. From left to right p=0.0007, 0.0995, 0.6903, and 0.7432, n=4, unpaired t-test. **(B)** Mouse BECs isolated from *Nfe2l2*^ENDO^ mice and *Nfe2l2*^Flox^ mice were cultured in vitro and TEER was measured for 24hrs. *p=0.0005, One-way ANOVA plus Sidak post hoc (n=8-9). **(C)** Scatterplot of RNA-seq analysis showing the influence of Nrf2 specific KO in ECs on BEC transcriptome at basal condition. Two weeks after tamoxifen-induced recombination, BECs were FCAS sorted from *Nfe2l2*^ENDO^ mice and their *Nfe2l2^fl/fl^* littermate’s control mice. Scatterplot was generated for genes with average expression >0.1 FPKM across the data sets. Highlighted with red and blue crosses are the genes whose expression are significantly increased or decreased respectively (*DESeq2 P*_adj<0.05, n=4-6). **(D)** Volcano plot of Mass Spectrometry analysis showing the influence of Nrf2 specific KO in ECs on BEC proteome at basal condition. Two weeks after tamoxifen-induced recombination, BECs were FCAS sorted from *Nfe2l2*^ENDO^ mice and their *Nfe2l2^fl/fl^* littermate’s control mice. Highlighted with red and blue dots are the proteins whose expression are significantly increased or decreased respectively (*p*<0.05, n=4).

### The Nrf2 activator RTA-404 modifies the BEC transcriptome under inflammatory conditions, which was abolished in endothelial cell-specific Nrf2 knockout mice

It is well established that systemic inflammation can lead to BBB compromise and neuroinflammation, so we employed a model of systemic inflammation to determine whether pharmacologically activating Nrf2 with RTA-404 can modify neuroinflammation. We induced systemic inflammation in mice by intraperitoneal injection of bacterial endotoxin LPS in *Nfe2l2*^LoxP/LoxP^ mice, chosen as a “conditional-ready” control to compare with conditional knockout mice used later in the study. BECs were sorted by FACS 24h post-LPS and RNA-seq performed, revealing a widespread inflammatory response in BECs, induced many pro-inflammatory genes, e.g. *Icam, Vcam, Ifitm1, Cxcl1, Ccl12, Csf3, Ccl5* and *Nfkbiz*, and down-regulated gene expression of tight junction proteins, e.g. *Cldn5 and Tjp1* (Fig. 3A). Pre-administration of RTA-404 significantly modulated the BEC transcriptome under conditions of LPS exposure (Fig. 3B). Known Nrf2 target genes, including *Gsta4, Gstp1, Gatm, Gamt* and *Aldh*, were induced. Gene expression of tight junction proteins including *Jam3*, *Gjc2* and *Gjc3* were significantly upregulated (*DESeq2 P*_adj<0.05; Fig.3B). Ingenuity Pathway Analysis (IPA) of genes significantly altered by RTA-404 revealed several down-regulated signatures associated with inflammation, particularly interferon signalling (Fig. 3C). In addition, mass spectrometry analysis showed that RTA-404 had a significant impact on the BEC proteome under conditions of LPS exposure, promoting expression of the proteins involved in tight junction function and ion transports across the membrane, e.g. Zo1, Slc6a6 and Slc38a5 (Fig. 3D). Thus, RTA-404 significantly modifies both BEC transcriptome and proteome following an inflammatory insult. We next wanted to investigate the site of action of RTA-404 with respect to its influence on the BEC transcriptome under conditions of LPS treatment, since the effect of RTA-404 could be due to direct activation of BEC Nrf2 or due to indirect effects of RTA-404 on the immune system response to LPS injection (intraperitoneal injection of LPS activates tissue-resident macrophages there as well as inducing inflammatory responses in circulating immune cells^4^). We then assessed the influence of RTA-404 on the BEC transcriptome of *Nfe2l2*^ENDO^ mice under conditions of LPS exposure. In contrast to the influence of RTA-404 on *Nfe2l2^fl/fl^* BECs, the effect of RTA-404 was essentially abolished in *Nfe2l2*^ENDO^ BECs (Fig. 3E). This strongly suggests that BEC Nrf2 is the locus of action for RTA-404, as opposed to upstream events such as LPS-induced immune cell activation.

**Fig.3.**
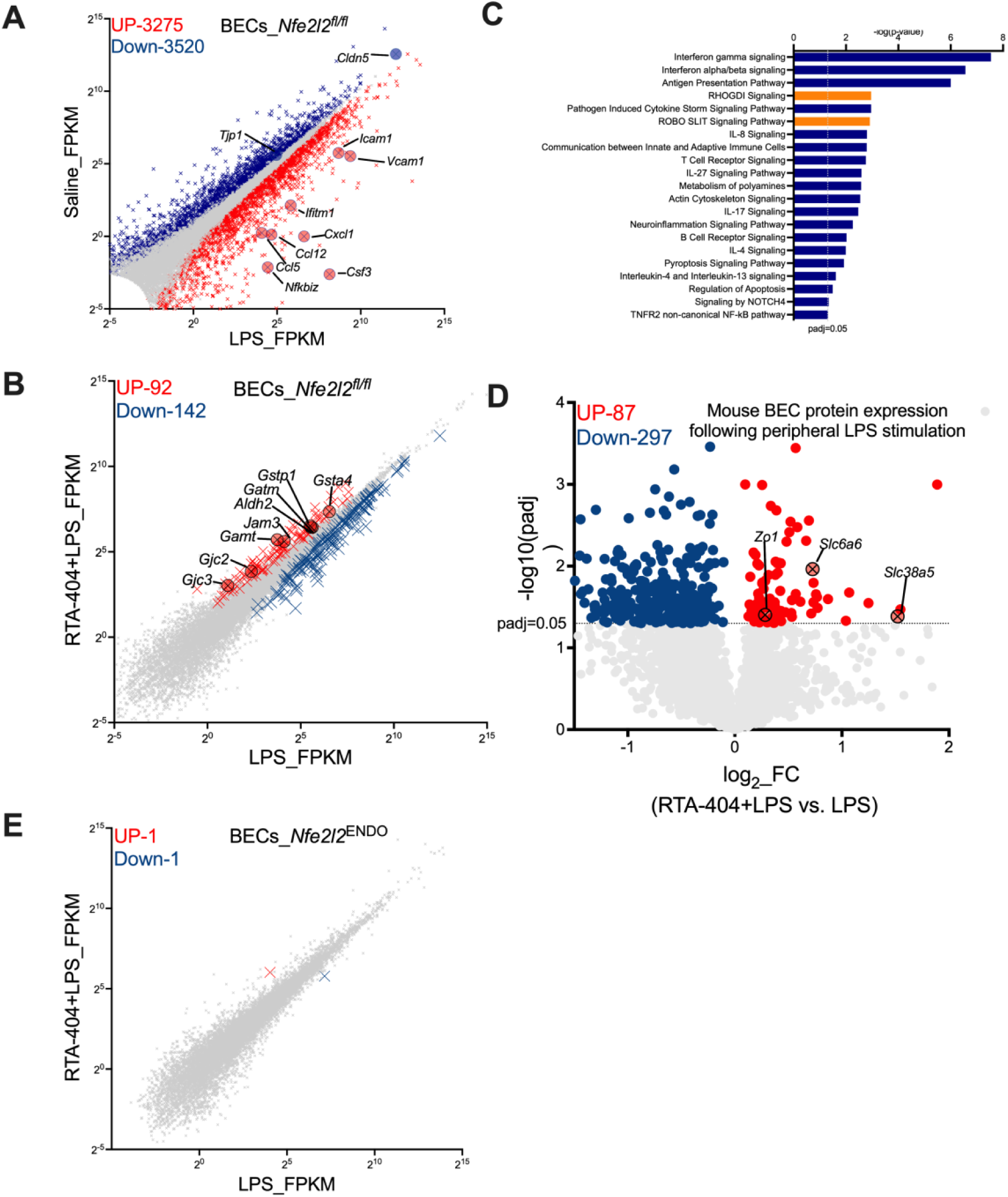
Nrf2 activator RTA-404 modifies the BEC transcriptome under inflammatory conditions. Mice were given i.p. injection of either saline or RTA-404 (n=6-8) daily for 4 days, following which either saline or LPS was given for 24 hrs. The brain cells were isolated and then sorted with FACS for BECs, RNA extracted and RNA-seq performed. Scatterplot was generated for genes with average expression >0.1 FPKM across the data sets. Highlighted with red and blue crosses are the genes whose expression are significantly increased or decreased respectively (*DESeq2 P*_adj<0.05, n=6-8). **(A)** Scatterplot of RNA-seq analysis showing the BEC transcriptome modified by peripheral LPS insult compared with saline in *Nfe2l2^fl/fl^* mice (LPS vs Saline). **(B)** Scatterplot of RNA-seq analysis showing the LPS-induced BEC transcriptome modified by pre-administration of RTA-404 in *Nfe2l2^fl/fl^* mice (LPS+RTA-404 vs. LPS+saline). **(C)** IPA analysis identifying activated or inhibited pathways by pre-administration of RTA-404 under conditions of LPS exposure (LPS+RTA-404 vs. LPS+saline). Significantly DEGs (*DESeq2 P*_adj<0.05, |Log_2_FC|>1, n=6-8) with BEC transcriptome as reference dataset were analyzed to calculate the p- value of overlap and z-score of overall activation/inhibition states of individual pathways. The significant activated (p<0.05, z-score >2) or inhibited (p<0.05, z-score<-2) pathways are shown in the bar chart in orange and blue respectively. **(D)** Volcano plot of Mass Spectrometry analysis showing the LPS-induced BEC proteome modified by pre-administration of RTA-404 in *Nfe2l2^fl/fl^* mice (LPS+RTA-404 vs. LPS+saline). **(E)** Scatterplot of RNA-seq analysis showing the LPS-induced BEC transcriptome modified by pre-administration of RTA-404 in *Nfe2l2*^ENDO^ mice.

### Endothelial cell Nrf2 maintains microglial homeostatic signature and regulates microglial activation under inflammatory conditions

Neuroinflammation following a peripheral inflammatory insult is characterised by microglial activation and reactive astrogliosis, and can involve infiltration of circulating immune cells, such as macrophages, into the brain parenchyma^2^. As expected, microglia and astrocytes displayed widespread transcriptional changes following LPS exposure (Fig. 4 A & Fig. 5A). Many proinflammatory genes, e.g. *Il1b, Msr1, Ccl5, ifitm3* and *Cxcl13* in microglia (Fig. 4A) and *Gfap*, *icam1, Ccl2, Cxl10* and *Ifitm1* in astrocytes (Fig. 5A) were induced. Moreover, microglial homeostatic genes, e.g. *Tmem119* and *P2ry12* were suppressed (Fig. 4A). Pre-administration of RTA-404 significantly modulated the transcriptome of microglia under conditions of LPS exposure (Fig. 4B). Of note, microglial homeostatic signature genes (MHSGs) such as *Tmem119*, *Sparc*, *Tgfbr1*, and *P2ry12* were significantly upregulated (Fig. 4B). To assess this systematically we took the set of MHSGs defined by Butovsky et al. (2014)^43^ and looked at the influence of pre-administration of RTA-404 on the expression of MHSGs under conditions of LPS exposure. Expression of these genes were significantly upregulated by pre-administration of RTA-404 under conditions of LPS exposure (Fig.4D), suggesting that pharmacologically activating Nrf2 has the capacity to maintain MHSG expression under inflammatory conditions. IPA analysis of genes significantly altered by RTA-404 in microglia revealed several down-regulated signatures associated with inflammation, particularly interleukin signalling, cell surface interactions at the vascular wall and leukocyte extravasation signalling (Fig. 4E). We performed immunohistochemistry to look at protein expression of microglia activation marker Iba1. RTA-404 significantly reduced Iba1 expression in response to peripheral LPS stimulation (Figure. S3. A). To further assess whether RTA-404 had the capacity to modulate microglial responses to LPS, we took microglial genes up- and down-regulated by LPS and studied the impact of RTA-404 pre-treatment. We found that the gene set induced by LPS in microglia was overall repressed by RTA-404 treatment (Fig. 4F (i & iii)), and the gene set repressed by LPS in microglia was overall elevated by RTA-404 treatment (Fig. 4G (i & iii)). Thus, RTA-404 was successful in suppressing microglial responses to peripheral LPS administration. We next asked whether the influence of RTA-404 on the microglial response involved activation of Nrf2 in ECs. Notably, when we performed the experiment on *Nfe2l2*^ENDO^ mice we found that the effect of RTA-404 on the microglial transcriptome was abolished (Fig. 4C). In addition, cross-reference analysis showed that no significant effect of RTA-404 was observed on LPS-induced changes in microglia (Fig. 4F (ii & iii) & Fig. 4G (ii & iii)). Thus, the impact of RTA-404 on microglia’s inflammatory response to peripheral LPS administration is mediated via BEC Nrf2 activation.

**Fig.4.**
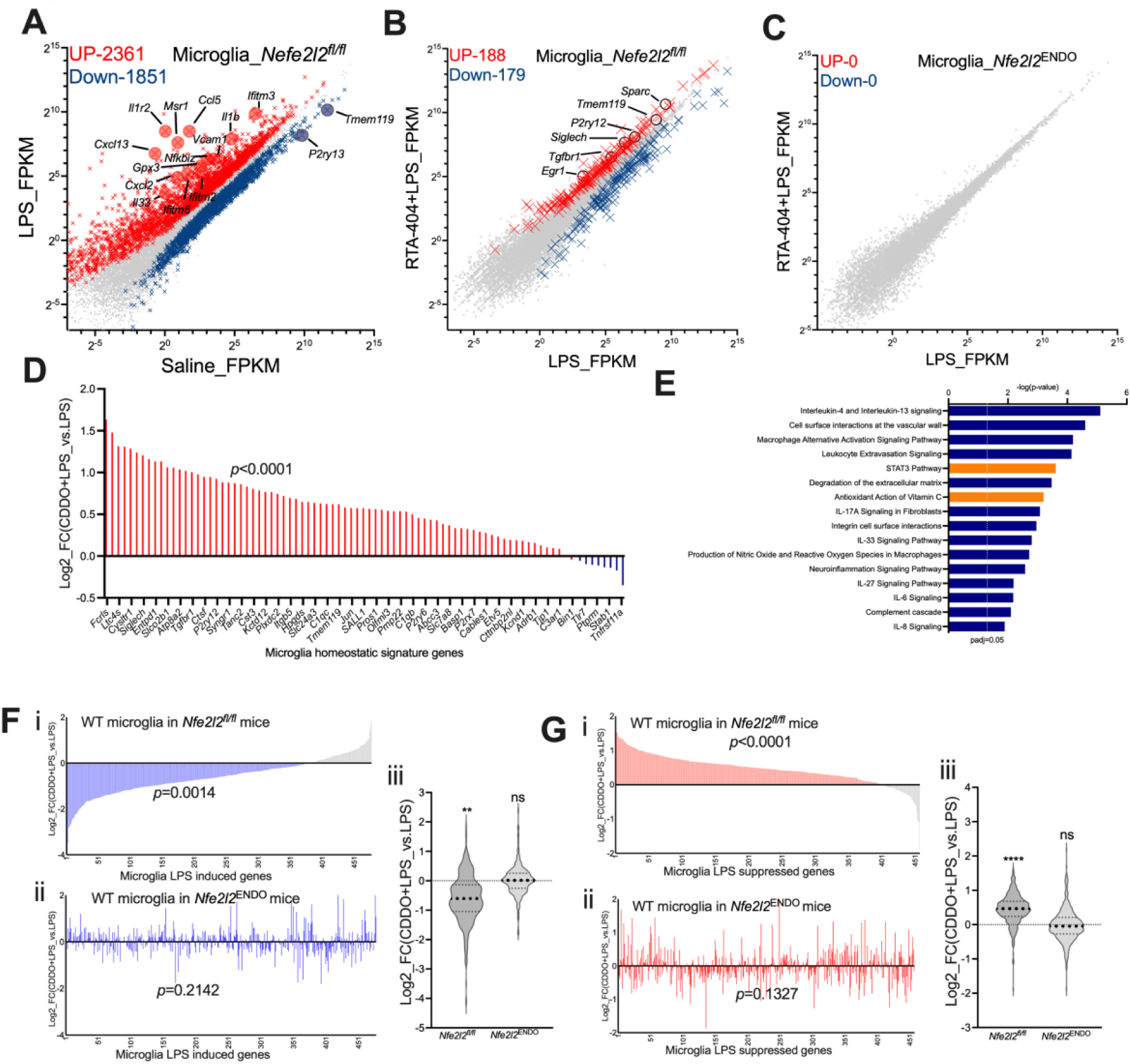
Endothelial cell Nrf2 maintains microglial homeostatic signature and regulates microglial activation under inflammatory conditions. Mice were given i.p. injection of either saline or RTA-404 (n=6-8) daily for 4 days, following which either saline or LPS was given for 24 hrs. The brain cells were isolated and then sorted with FACS for microglia, RNA extracted and RNA-seq performed. Scatterplot was generated for genes with average expression >0.1 FPKM across the data sets. Highlighted with red and blue crosses are the genes whose expression are significantly increased or decreased respectively (*DESeq2 P*_adj<0.05, n=6-8). **(A)** Scatterplot of RNA-seq analysis showing the microglia transcriptome modified by peripheral LPS insult compared with saline in *Nfe2l2^fl/fl^* mice (LPS vs Saline). **(B)** Scatterplot of RNA-seq analysis showing the LPS-induced microglia transcriptome modified by pre-administration of RTA-404 in *Nfe2l2^fl/fl^* mice (LPS+RTA-404 vs. LPS+saline). **(C)** Scatterplot of RNA-seq analysis showing the LPS-induced microglia transcriptome modified by pre-administration of RTA-404 in *Nfe2l2*^ENDO^ mice. **(D)** RTA-404 promotes microglial homeostatic signature under inflammatory conditions. Genes considered are those expressed >0.5 FPKM in our data and within the group of microglia signature genes defined by (Butovsky et al., 2014). For each gene, log_2_ fold change (log_2_FC) of microglia gene expression in *Nfe2l2^fl/fl^* mice by pre-administration of RTA-404 under conditions of LPS exposure (LPS+RTA-404 vs. LPS+saline) is shown. The data were mined from the complete set shown in (B). *P<0.0001, F(1, 498)= 95.1 relates to main effect of pre-administration of RTA-404 on microglia signature gene expression under inflammatory conditions, two-way ANOVA. **(E)** IPA analysis identifying activated or inhibited pathways in microglia by pre-administration of RTA- 404 under conditions of LPS exposure (LPS+RTA-404 vs. LPS+saline). Significantly DEGs (*DESeq2 P*_adj<0.05, |Log_2_FC|>1, n=6-8) with microglia transcriptome as reference dataset were analyzed to calculate the p-value of overlap and z-score of overall activation/inhibition states of individual pathways. The significant activated (p<0.05, z-score >2) or inhibited (p<0.05, z-score <-2) pathways are shown in the bar chart in orange and blue respectively. **(F)** The influence of RTA-404 on LPS-induced genes in microglia. For LPS-induced genes (x-axis, FPKM>1*, DESeqP*_adj<0.05, Log_2_FC>1) in microglia in *Nfe2l2^fl/fl^* mice, the Log_2_FC (y-axis) of each gene following treatment with LPS ± RTA-404 in microglia in either *Nfe2l2^fl/fl^* mice or *Nfe2l2*^ENDO^ mice were plotted. (iii) *p=0.0014, F (1,6720) =10.24 relates to main effect of RTA-404 on microglial gene sets induced by LPS, 2-way ANOVA. **(G)** The influence of RTA-404 on LPS-repressed genes in microglia. For LPS-repressed genes (x-axis, FPKM>1*, DESeqP*_adj<0.05, Log_2_FC <-1) in microglia in *Nfe2l2^fl/fl^* mice, the log2 fold change (y-axis) of each gene following treatment with LPS ± RTA-404 in microglia in either *Nfe2l2^fl/fl^* mice or *Nfe2l2*^ENDO^ mice were plotted. (iii): *P<0.0001, F (1,2072) =20.12 relates to main effect of RTA-404 on microglial gene sets suppressed by LPS, 2-way ANOVA.

### Endothelial cell Nrf2 regulates astrogliosis under inflammatory conditions

We next analysed astrocytes sorted from LPS-treated *Nfe2l2^fl/fl^* and *Nfe2l2*^ENDO^ mice. As with microglia, pre-administration of RTA-404 significantly modulated the transcriptome of astrocytes under conditions of LPS exposure (Fig. 5B). Of note, the expression of classic astrogliosis marker *Gfap* was downregulated by RTA-404 under LPS exposure (Fig. 5B), although immunohistochemistry did not detect significant change in Gfap protein expression by RTA-404 (Figure. S3. B). IPA pathway analysis shows of genes significantly altered by RTA-404 in astrocytes revealed down-regulated signatures associated with inflammation, particularly interferon signalling and interleukin signalling (Fig. 5D). Like in microglia, we observed an RTA-404-induced suppression of LPS-induced inflammatory responses in astrocytes: RTA-404 significantly inhibited the set of LPS-activated genes (Fig. 5E (i & iii)), and up-regulated the set of LPS-repressed genes (Fig. 5F (i & iii)). Moreover, the effect of RTA-404 on the astrocyte transcriptome under conditions of LPS exposure was abolished in *Nfe2l2*^ENDO^ mice (Fig. 5E) and no significant effect of RTA-404 was observed on LPS-induced changes in astrocytes (Fig. 5E (ii & iii) & Fig. 5F (ii & iii)). Thus, despite the potential of RTA-404 to activate Nrf2 directly in microglia and astrocytes, the effects of RTA-404 on these cell types are in fact dependent on Nrf2 in endothelial cells.

**Fig.5.**
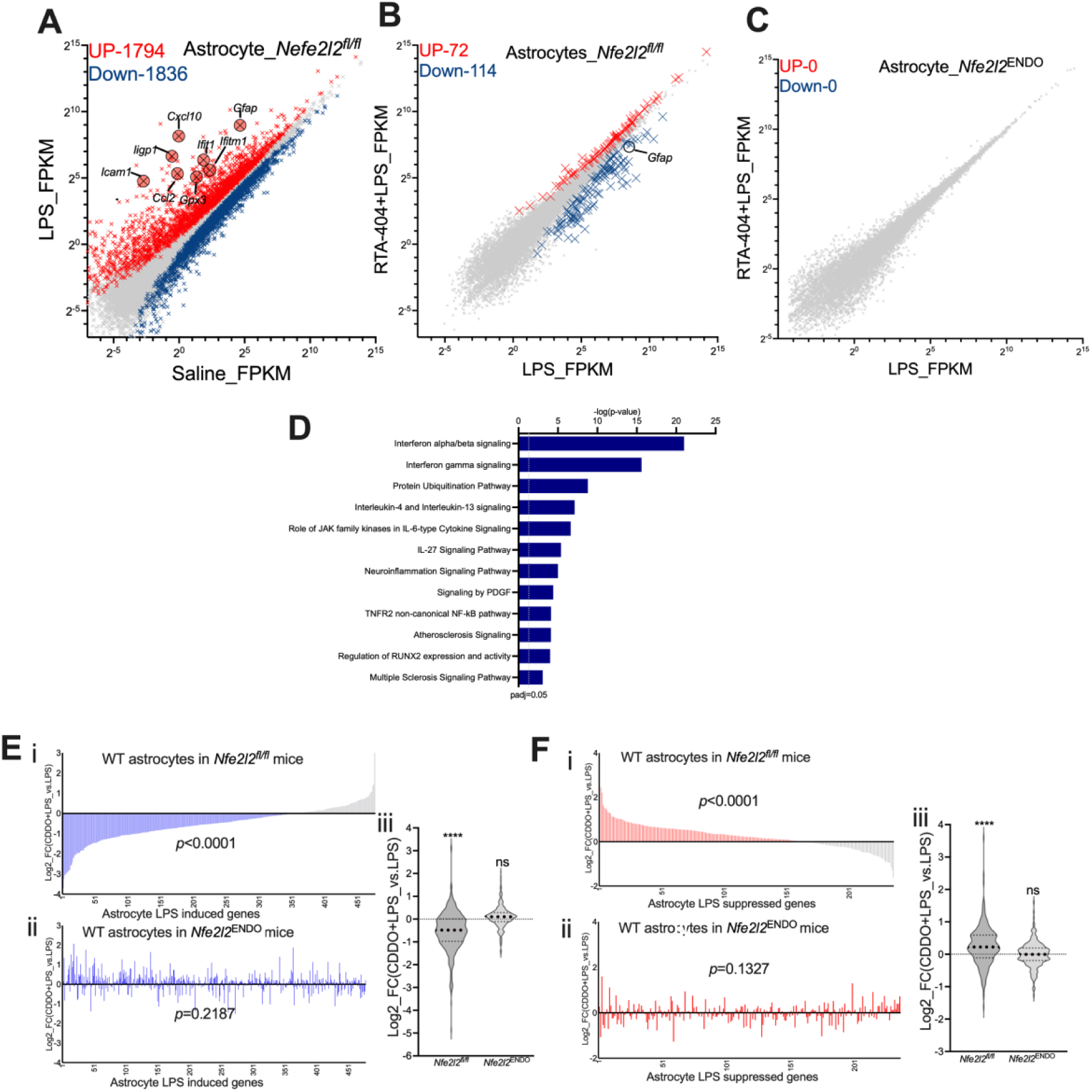
Endothelial cell Nrf2 is a key point of regulating astrogliosis under inflammatory conditions. Mice were given i.p. injection of either saline or RTA-404 (n=6-8) daily for 4 days, following which either saline or LPS was given for 24 hrs. The brain cells were isolated and then sorted with FACS for astrocytes, RNA extracted and RNA-seq performed. Scatterplot was generated for genes with average expression >0.1 FPKM across the data sets. Highlighted with red and blue crosses are the genes whose expression are significantly increased or decreased respectively (*DESeq2 P*_adj<0.05, n=6-8). **(A)** Scatterplot of RNA-seq analysis showing the astrocyte transcriptome modified by peripheral LPS insult compared with saline in *Nfe2l2^fl/fl^* mice (LPS vs Saline). **(B)** Scatterplot of RNA-seq analysis showing the LPS-induced astrocyte transcriptome modified by pre-administration of RTA-404 in *Nfe2l2^fl/fl^* mice (LPS+RTA-404 vs. LPS+saline). **(C)** Scatterplot of RNA-seq analysis showing the LPS-induced astrocyte transcriptome modified by pre-administration of RTA-404 in *Nfe2l2*^ENDO^ mice. **(D)** IPA analysis identifying activated or inhibited pathways in astrocytes by pre-administration of RTA- 404 under conditions of LPS exposure (LPS+RTA-404 vs. LPS+saline). Significantly DEGs (*DESeq2 P*_adj<0.05, |Log_2_FC|>1, n=6-8) with astrocyte transcriptome as reference dataset were analyzed to calculate the p-value of overlap and z-score of overall activation/inhibition states of individual pathways. The significantly inhibited (p<0.05, z-score <-2) pathways are shown in the bar chart in blue. **(E)** The influence of RTA-404 on LPS-induced genes in astrocytes. For LPS-induced genes (x-axis, FPKM>1*, DESeqP*_adj<0.05, Log_2_FC >1) in astrocytes in *Nfe2l2^fl/fl^* mice, the Log_2_FC (y-axis) of each gene following treatment with LPS ± RTA-404 in astrocytes in either *Nfe2l2^fl/fl^* mice or *Nfe2l2*^ENDO^ mice were plotted. (iii) *p<0.0001, F (1,6056) =29.29 relates to main effect of RTA-404 on astrocytic gene sets induced by LPS, 2-way ANOVA. **(F)** The influence of RTA-404 on LPS-repressed genes in astrocytes. For LPS-repressed genes (x-axis, FPKM>1*, DESeqP*_adj<0.05, Log_2_FC <-1) in astrocytes in *Nfe2l2^fl/fl^* mice, the Log_2_FC (y-axis) of each gene following treatment with LPS ± RTA-404 in astrocytes in either *Nfe2l2^fl/fl^* mice or *Nfe2l2*^ENDO^ mice were plotted. (iii): *p<0.0001, F (1,25556) =68.12 relates to main effect of RTA-404 on astrocytic gene sets suppressed by LPS, 2-way ANOVA.

### Endothelial cell Nrf2 regulates immune cell brain infiltration under inflammatory conditions

Neuroinflammation due to a peripheral insult can involve infiltration of circulating immune cells via a compromised BBB^2^. Since our observation that RTA-404 strengthened the TEER of a BEC monolayer in vitro, we wanted to assess whether RTA-404 could prevent BBB compromise. In our model of i.p. LPS administration we observe substantial peripheral macrophage infiltration into the brain parenchyma, assessed by flow cytometry, where CD11b^+^CD45^low^ microglia and CD11b^+^CD45^high^ macrophages can be distinguished. To confirm the gating strategy for brain microglia and macrophage populations in flow cytometry analysis, we took advantage of *Csf1r*^ΔFIRE/ΔFIRE^ mice in which brain microglia are lacking but other macrophage populations are unaffected (Fig. S1B). We observed parenchymal infiltration of CD11b^+^CD45^high^ macrophages upon peripheral LPS administration (Fig. 6A (i & ii) and Fig.6B) in *Nfe2l2^fl/fl^* mice. Importantly, RTA-404 administration abolished macrophage infiltration into the brain (Fig. 6A (iii) and Fig. 6B). Strikingly, the effect of RTA-404 in preventing macrophage infiltration into the parenchyma was abolished in *Nfe2l2*^ENDO^ mice (Fig.6A (vi) and Fig.6B). Thus, EC Nrf2 can play a key role in preventing immune cell infiltration into the brain parenchyma following peripheral inflammation.

**Fig.6.**
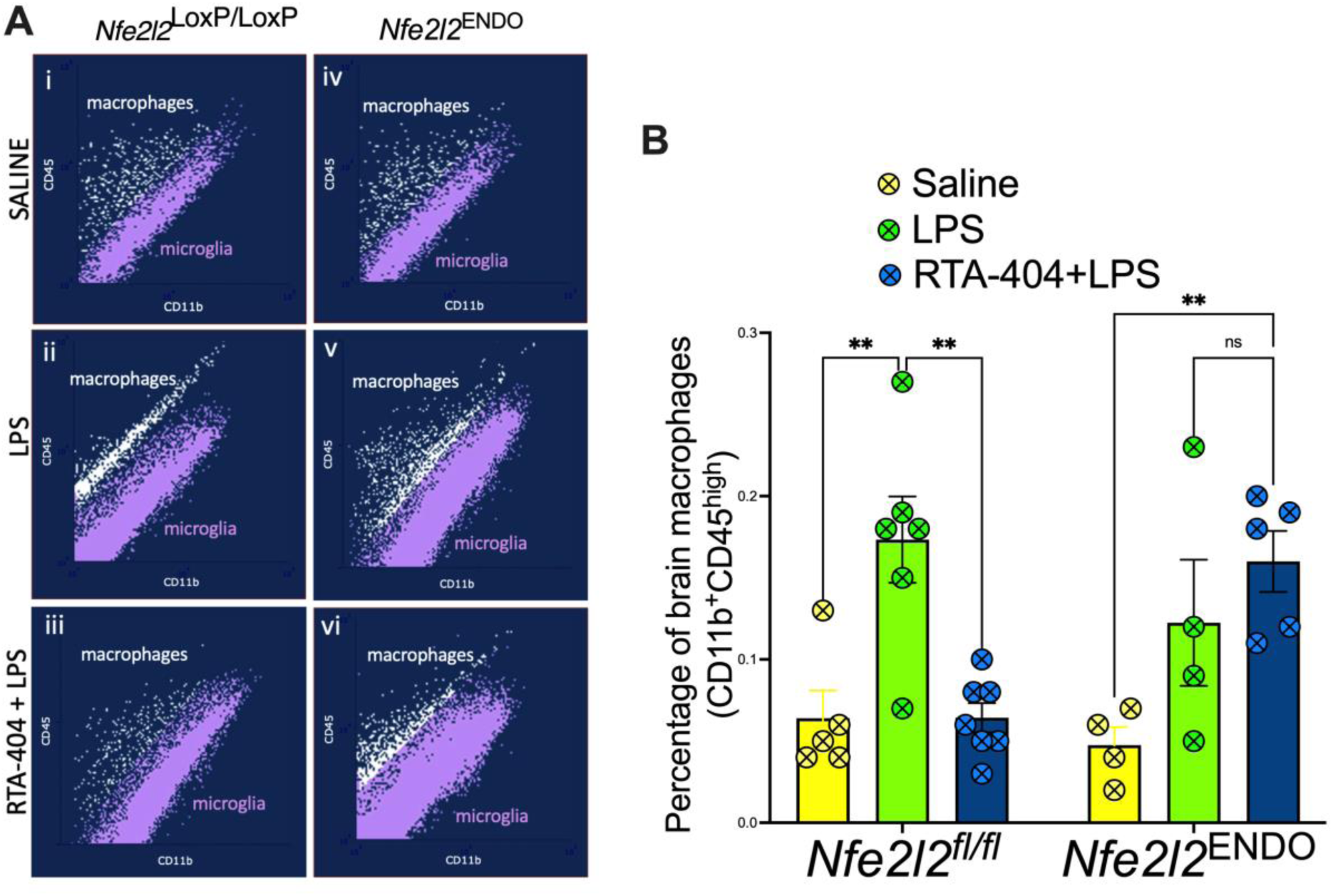
Endothelial cell Nrf2 regulates immune cell brain infiltration under inflammatory conditions. Mice were given i.p. injection of either saline or RTA-404 (n=6-8) daily for 4 days, following which either saline or LPS was given for 24 hrs. The brain cells were analyzed using flow cytometer and the number of macrophages (CD11b^+^CD45^high^) in the brain following LPS or RTA-404 treatment were quantified. **(A)** Representative flow cytometry graphs showing microglia and macrophage population in the brain. **(B)** Statistical analysis showing the effect of either LPS or RTA-404 on peripheral monocyte brain infiltration. p=0.0023, 0.0010, 0.0045,0.4774 from left to right, 2-way ANOVA plus Sidak post hoc (n=6- 8).

## Discussion

Our current study using cultured human BECs and inducible conditional EC-specific Nrf2 KO mice demonstrated that Nrf2 signalling plays a key role in maintaining BEC homoeostasis and BEC barrier strength through modifying BEC transcriptome and proteome. In addition to activating the classic Nrf2/HO antioxidant/detoxification pathway and promoting the expression of tight junction and adherent junction proteins, our current study identified several other genes and signalling pathways regulated by RTA-404 in BECs as the potential mechanisms that might contribute to promoting BEC barrier strength. Previous studies have suggested that synthetic oleanane triterpenoids including RTA-404 are multi-functional drugs, in addition to KEAP1/Nrf2/ARE pathway as a primary target, they can also interact with other pathways, e.g. IKK/NfκB, JAK/STAT, and PI3K/Akt^31,44^. Indeed, our current study also found multiple other pathways, e.g. IL10 signalling, iNOS signalling, and Insulin Receptor Signalling, Oxidative Phosphorylation, Cell Cycle Checkpoint were regulated by RTA-404 in human BECs at basal conditions, whether they were directly modified by RTA-404 or indirectly via Nrf2 pathway requires further investigation. Our current study also found that Nrf2 activation in ECs regulates immune system-to-brain communication in response to peripheral infection in mice. The mechanisms by which Nrf2 activation suppresses neuroinflammation may involve a combination of supressing EC inflammatory responses (e.g. EC chemokine release) as well as maintaining BBB integrity to prevent immune cell infiltration into the brain parenchyma. As such, BEC Nrf2 may play a classical anti-inflammatory role, but also a cell-type specific roles such as regulation of barrier integrity. Indeed, tight junction protein Jam3^45^ and Zo1 are up-regulated by RTA-404 treatment in BECs and down-regulated by Nrf2 KO in BECs. However, further investigations are needed to define the precise mechanism behind EC Nrf2-dependent prevention of immune cell infiltration.

As well as providing a greater understanding of the cell type-specific roles of Nrf2, our work suggests that EC Nrf2 is a tractable therapeutic target for neuroinflammation, with the advantage of being a CNS-relevant target that can be targeted by non-CNS-penetrating drugs. The growing appreciation of the long-lasting impact of immune system-to-brain signalling on cognitive function and neurodegenerative disease trajectory makes it essential to develop strategies to combat this^46,47^. The positive impact of RTA-404 in limiting neuroinflammation, coupled with the recent clinical success of its close relative RTA-408, suggest that administration of these or similar compounds may limit the CNS-adverse effects of peripheral inflammation. The effect of RTA-404 is striking, as is the fact that its effects are abolished in endothelial cell-specific Nrf2 deficient mice. We cannot rule out that RTA-404 is influencing events upstream of BECs (peripheral ECs) or downstream events (e.g. microglial and astrocyte responses). However, the striking abolition of the effects of RTA-404 in *Nfe2l2*^ENDO^ mice point to EC Nrf2 being necessary. We conclude that EC Nrf2 acts as a gatekeeper of BBB integrity and neuroinflammation and is potentially a promising therapeutic target for preventing systemic inflammation induced neuroinflammation in a range of clinical settings.

### Limitations of the study

One limitation of this study is that our genetic strategy employed to knock-out NRF2 does so in all ECs. An alternative strategy would be needed to be knock out NRF2 in BECs only in order to definitively implicate BEC NRF2 as a gatekeeper of neuroinflammation. While a mouse line has been generated that expresses Cre-ERT2 in BECs and no other ECs (*Slco1c1-*CreER^T2^ mice^28^) it also is expressed in a proportion of astrocytes^28^ which would be problematic for our study, given the high level of expression and functional importance of astrocytic NRF2^31^. The other limitation of the study is that while we show EC NRF2 to be *necessary* for the effect of RTA-404 on neuroinflammation, we don’t show that EC NRF2 is sufficient to prevent neuroinflammation following a peripheral insult. This would likely require EC or BEC-specific over-expression of NRF2 by generating a new transgenic mouse or EC-targeted AAVs (such as AAV9-X1^30^). Such gain-of-function experiments are part of our ongoing plans.

## STAR★Methods

### Key Resources Table

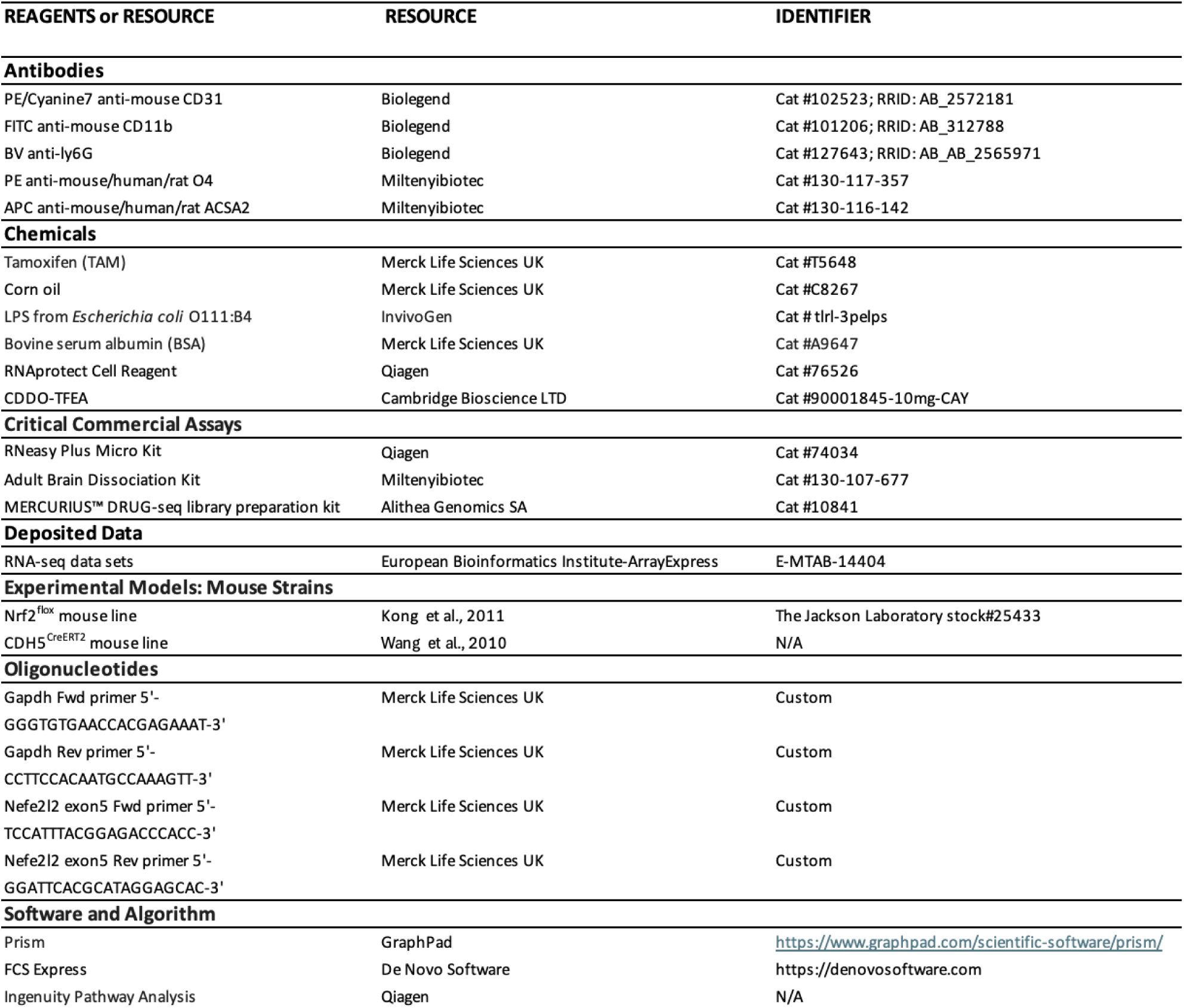

## Resource availability

### Lead Contact

Further information and requests for resources and reagents should be directed to and will be fulfilled by the Lead Contact, Jing Qiu.

### Materials Availability

This study did not generate new unique reagents.

### Data and Code Availability

All RNA-seq data that support the findings of this study is available at the European Bioinformatics Institute (ArrayExpress: E-MTAB-14404). All other data are available from the lead contact upon reasonable request.

### Experimental Model and Subject Details

#### Mice

The animals involved in this study were all on C57BL/6 background. The CDH5^CreERT2^ mouse line was originally developed in the laboratory of Dr. Ralf Adams^28^ and kindly provided by Prof. Andy Baker of University of Edinburgh. The Nrf2^flox^ mouse line was originally generated in the laboratory of Dr. Shyam Biswal and imported from Jackson Laboratories (Mouse Strain Number – 025433). The Nrf2^flox^ mice carrying *loxP* sites flanking exon 5 of the *Nfe2l2* gene were crossed to the CDH5^CreERT^^2^ mouse line^28^ to generate *Cdh5^CreERT^*^2^*: Nefe2l2^fl/fl^* mice. Male mice have been used throughout the study. All procedures described were performed at the University of Edinburgh in compliance with the UK Animals (Scientific Procedures) Act 1986 and University of Edinburgh regulations and carried out under project license numbers P2262369. Mice were group-housed in environmentally enriched cages within humidity and temperature-controlled rooms, with a 12-h light dark cycle with free access to food and water. Mouse genotypes were determined using real-time PCR with transgene specific probes (Transnetyx, Cordova, TN) unless otherwise stated.

## Method Details

### Tamoxifen, RTA-404 and LPS Treatments

For induction of Cre recombinase activity, 6-8-week-old *Cdh5^CreERT^*^2^*: Nefe2l2^fl/fl^* mice were given orally with 8 mg tamoxifen (TAM, T5648, Sigma-Aldrich, UK) solved in corn oil (C8267, Sigma-Aldrich, UK) at 4 time points with 24hrs apart. For all experiments, littermates carrying the respective loxP-flanked alles but lacking expression of Cre recombinase (+/+ TAM) were used as controls. Male mice have been used throughout the study. The *Cdh5^CreERT^*^2^*: Nefe2l2^fl/fl^* mice and control mice were injected peritoneally with 5mg/kg of RTA-404 (90001845-10mg-CAY, CAMBRIDGE BIOSCIENCE LTD, UK) for 4 consecutive days with 24 hrs apart, following which 3mg/kg of LPS (tlrl-3pelps, Invivogen, US) was given intraperitoneally to the mice for 24 hrs.

### Isolation of brain cells, Cell Sorting, and RNA extraction

Single brain cells were isolated using Adult Brain Dissociation kit (Miltenyie, 130-107-677, Germany) as per the manufacturer’s instructions. The single brain cell suspension was then incubated with antibodies against CD31 (Biolegend, 102523, UK) CD11b (Biolegend, 101205, UK), ACSA2 (Miltenyie, 130-116-245, Germany), O4 (Miltenyie, 130-117-357, Germany), CD45 (Biolegend, 103125, Germany) for 15 minutes in ice, washed with PBS and then immediately FACS sorted into RNAprotect Cell Reagent (Qiagen, 76526, UK) under the gates of CD31^+^CD45^-^ ACSA2^-^ for BECs, LY6G^-^CD11b^+^CD45^low^ for microglia, ACSA^+^O4^-^ for astrocytes and O4^+^ for oligodendrocytes (Fig. S1). RNA extraction was carried out using the RNeasy Plus Micro Kit (Qiagen) as per the manufacturer’s instructions.

### Mass spectrometry

The FACs sorted BECs were denatured with 6M guanidine hydrochloride and then treated with 5mM tris (2-carboxyethyl) phosphine (TCEP) and 10mM chloroacetaldehyde (CAA) for reduction and alkylation respectively, before heating at 95°C for 5 minutes. After cooling to room temperature, the samples were digested first with 0.2 μg endoproteinase LysC in 3M guanidine at 37°C overnight and then with 0.3 μg trypsin in 1M guanidine at 37°C for 4 hours. The resulting peptides were cleaned up using C18 Stage Tips (Rappsilber et al., 2003), separated with an Ultimate 3000-series RSLC Nano System (Thermo Fisher), ionised with an IonOpticks Aurora C18 nano packed emitter in a Proxeon nano source (Thermo Fisher), analyzed on a Orbitrap Fusion Lumos Tribrid Mass Spectrometer (Thermo Fisher), and identified and counted using MaxQuant platform (version 1.6.7.0) and Uniprot proteomics database (Cox and Mann, 2008; UniProt Consortium, 2019).

### Immunohistochemistry

Primary antibodies are listed in the table below. All counts and analyses were performed rostral to Bregma in the forebrain of the mice as defined previously^48,49^. Three slices per animal were used (and combined to create a single average). For imaging of Iba1 and GFAP slices, animals were killed by perfusion-fixation with 4% PFA. Brains were removed and 30 μM slices prepared using a cryostat. Slices were washed 3x with PBS, blocked and permeabilized with 10% normal goat serum (Thermofisher) with 0.3% Triton-X (Sigma-Aldrich). Primary antibodies were applied overnight at 4 °C whilst shaking, and secondary antibodies applied for 2 hours at RT. Following this, cells or slices were washed 4× in PBS and mounted using Vectashield with 4′,6-diamidino-2-phenylindole (DAPI) (Vector Labs). Images were acquired using a Nikon A1R laser scanning confocal microscope. Laser beams of wavelength 488 and 561 nm were used to excite FITC and CY3 fluorophores respectively, using a 20 × 0.8 NA objective lens. NIS elements software was used for the image acquisition and ImageJ software was used for the analysis. For immunohistochemistry in cultures cells, a paraformaldehyde fixation followed by permeabilization (NP40)^52^.

### Human BECs culture and TEER measurement

Human Brain Microvascular Endothelia Cells (BECs) were purchase from Innoprot (P10361-IM, Elexalde Derio, Spain). All plates and dishes for endothelial cells culture were coated with 0.2% Gelatine (G9391, Sigma-Aldrich) and 10ug/mL fibronectin (F1141-5MG, Sigma-Aldrich) for 30 min at 37 °C. Cells were cultured in Endothelial Cell Growth Medium with premixed supplements (C22022, Promocell) and 1% antibiotic-antimycotic solution (11570486, Gibco). For TEER measurement, the Electric Cell-substrate Impedance Sensing (ECIS) machine were used, implemented with 8-well 8W10E+ arrays via a ECIS Z-Theta station (Applied Biophysics). Cells were seeded into each well of the pre-coated array and allowed to adhere overnight. When the cells were confluent, 400 nM RTA-404 were added in the medium and the TEER recorded at a frequency of 4 kHz for 24 hours.

### Quantitative RT-PCR

cDNA was generated using the Transcriptor First Strand cDNA Synthesis Kit (Roche). Seven microlitres of RNA was added to the RT and buffer mixture prepared with random hexamers and oligoDT primers as per kit instructions, and qRT-PCR carried out with the following programme: 10 min at 25 °C, 30 min at 55 °C and 5 min at 85 °C. qPCRs were run on a Stratagene Mx3000P QPCR System (Agilent Technologies) using SYBR Green MasterRox (Roche) with 6 ng of cDNA per well of a 96-well plate, using the following programme: 10 min at 95 °C, 40 cycles of 30 s at 95 °C, 40 s at 60 °C and 30 s at 72 °C, with a subsequent cycle of 1 min at 95 °C and 30 s at 55 °C ramping up to 95 °C over 30 s (to measure the dissociation curve).

### RNA-seq and Analysis

For standard TruSeq (Experiments presented in Figures 2A, 3B, 4A, and 5A), libraries were prepared by Edinburgh Genomics using the Illumina TruSeq stranded mRNA-seq kit, according to the manufacturer’s protocol (Illumina). The libraries were pooled and sequenced to 75 base paired-end on an Illumina NovaSeqTM 6000 to a depth of approximately 50 million paired-end reads per sample. For barcoded mRNA sequencing (Experiments presented in Figures 2B, 3C, 4B & C, 5B & C), RNA sequencing libraries were prepared using the MERCURIUS Drug-seq v2 library Preparation 96 kit as per the manufacturer’s instructions. Briefly, each RNA sample was reverse transcribed in a 96-well plate with individual barcoded oligo-dT primers. Next, all the samples were pooled together, purified using the DNA Clean and Concentrator kit (Zymo Research, Cat.#D4014), and treated with exonuclease I. Double-stranded cDNA was generated by the second stand synthesis via the nick translation method. Full-length double-stranded cDNA was purified with 30 μL (0.6x) of AMPure XP magnetic beads (Beckman Coulter, Cat.#A63881) and eluted in 20 μL of water. The Illumina compatible libraries were prepared by tagmentation of 5 ng of full-length double-stranded cDNA with 1 μL of in-house produced Tn5 enzyme (11 μM). The final library was amplified using 15 cycles and the fragments ranging 200–1000 bp were size-selected using AMPure beads (Beckman Coulter, Cat.#A63881) (first round 0.5x beads, second 0.7x). The libraries were profiled with the High Sensitivity NGS Fragment Analysis Kit (Advanced Analytical, Cat.#DNF- 474) and measured with the Qubit dsDNA HS Assay Kit (Invitrogen, Cat.#Q32851) prior to pooling and sequencing using the Illumina NextSeq 2000 P2 platform. For read mapping and feature counting, genome sequences and gene annotations were downloaded from Ensembl version 94. Differential expression (DGE) analysis on data sets was performed using DESeq2 (R package version 1.18.1) using a significance threshold set at a Benjamini-Hochberg-adjusted p-value of 0.05.

## Acknowledgements

We would like to thank Prof. Andrew Baker for providing the CDH5^CreERT2^ mouse Line.

We would like to thank the Flow Cytometry Facility, led by Dr. Fiona Rossi, at the Centre for Regenerative Medicine at the University of Edinburgh for their help with the Flow Cytometry work.

We would like to thank Prof. Alex Von Kriegsheim’s group at the Centre for Cancer Research at the University of Edinburgh for performing the Mass Spectrometry analysis.

The research leading to these results received funding from the Ann Rowling Regenerative Neurology Clinic and the UK DRI at the University of Edinburgh.

## Author contributions

H.Z., Z.H. and T.L. performed the experiments. X.H., O.D., H.Z., and J.Q. analysed the data. J.Q. conceived the study and wrote the manuscript.

---

**Fig. S1.**
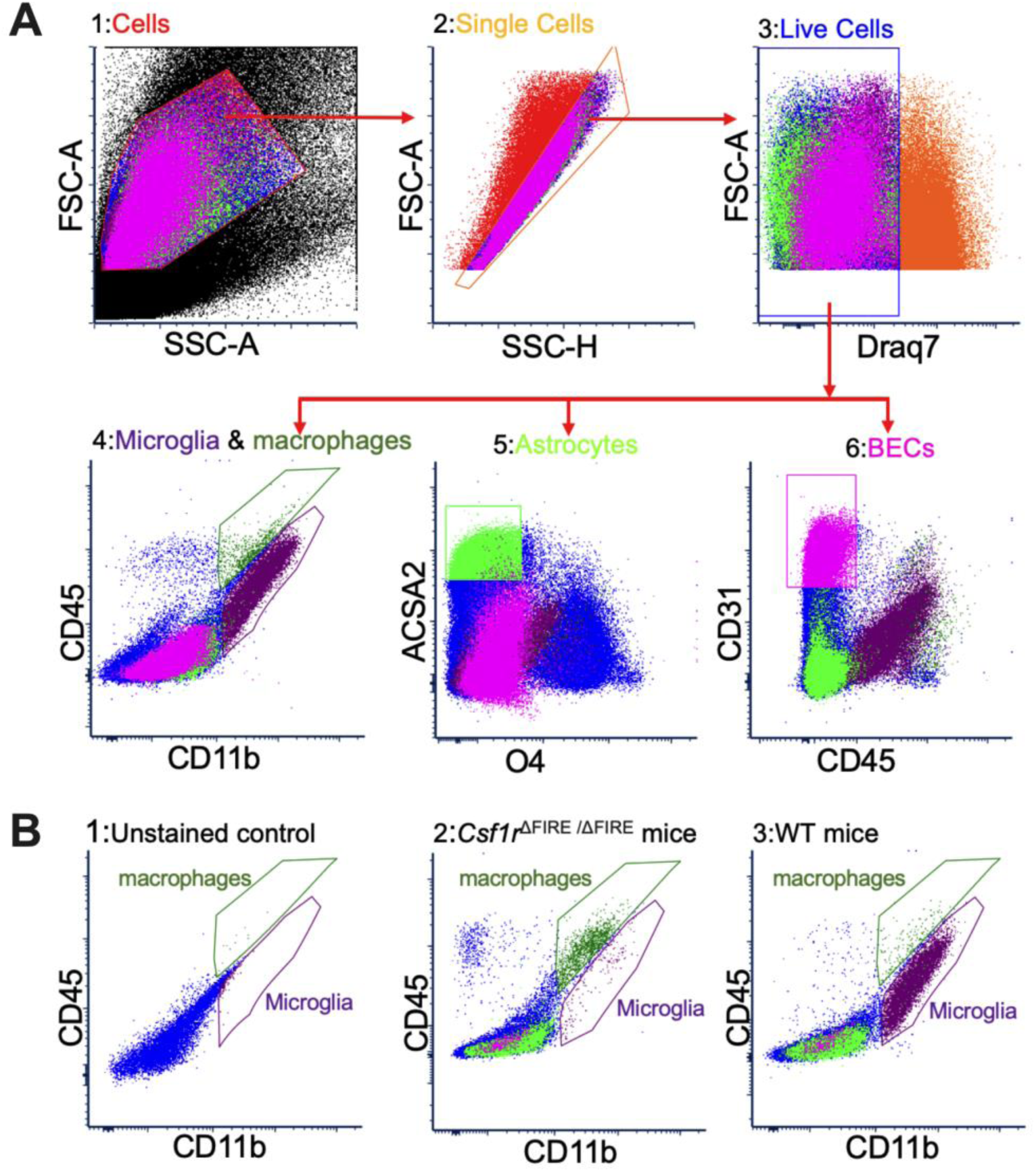
FACS gating strategy for BECs, astrocytes, microglia and macrophages. **(A)** Representative flow cytometry graphs are shown for gating strategy for populations of BECs (CD31^+^CD45^-^), astrocytes (ACSA2^+^O4^-^), microglia (CD11b^+^CD45^low^) and macrophages (CD11b^+^CD45^high^). Brain cells were isolated as single cell suspension from WT mice and labelled with full panel of antibodies for BECs, astrocytes, microglia and macrophages. **(B)** Representative flow cytometry graphs are shown for establishing gating strategy for microglia (CD11b^+^CD45^low^) and macrophages (CD11b^+^CD45^high^). 1. Brain cells from WT mice without antibody labelling. 2 & 3. Brain cells from *Csf1r*^ΔFIRE/ΔFIRE^mice (2) and WT mice (3) labelled with a panel of antibodies for BECs, astrocytes, microglia and macrophages.

**Fig. S2.**
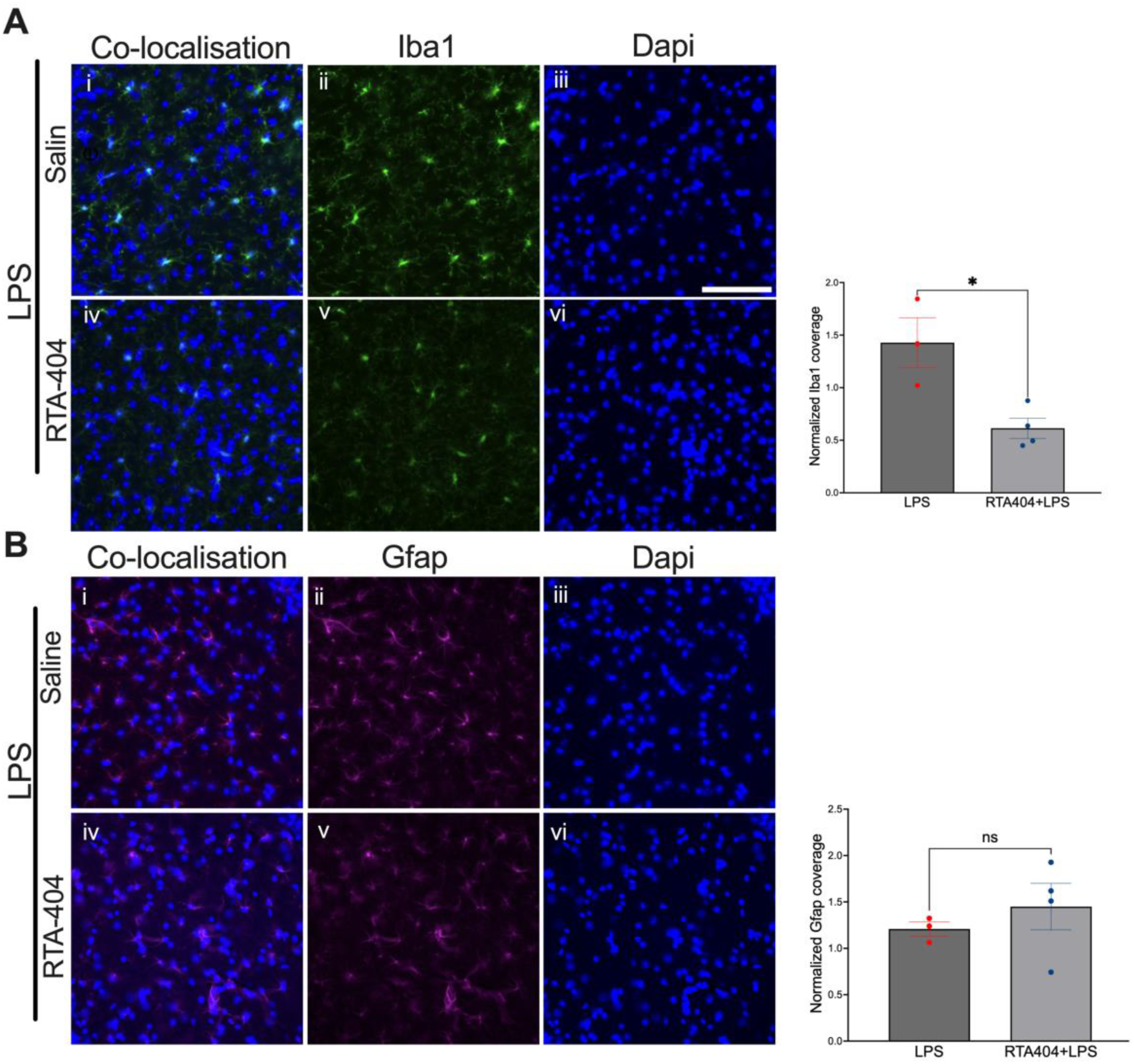
The influence of RTA-404 on microglia and astrocytes protein expression. **(A)** Immunohistochemistry showing the influence of RTA-404 on Iba1 protein expression in microglia in *Nfe2l2^fl/fl^* mice. p=0.0164, unpaired t-test. **(B)** Immunohistochemistry showing the influence of RTA-404 on Gfap protein expression in astrocyte in *Nfe2l2^fl/fl^* mice. P=0.4637, unpaired t-test.

**Table S1.**
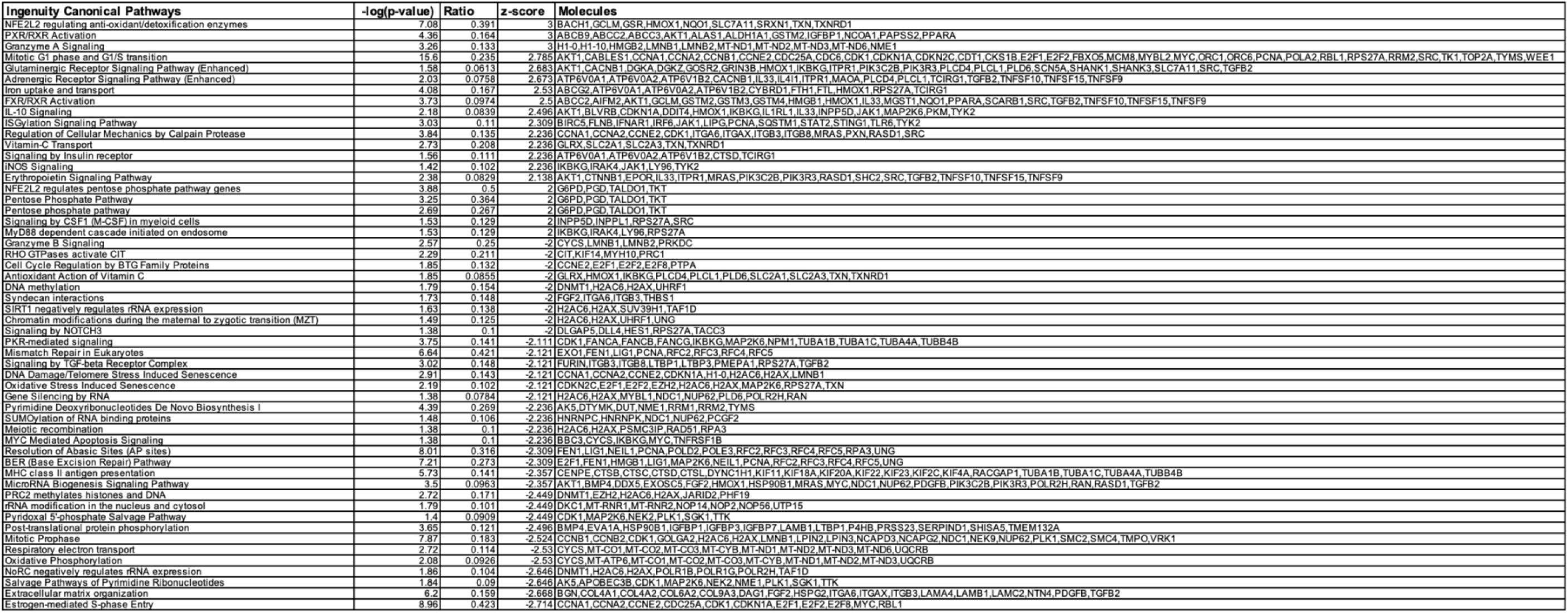

